# Characterization of the bioaccumulation and toxicity of copper pyrithione, an antifouling compound, on juveniles of rainbow trout

**DOI:** 10.1101/2023.01.31.526498

**Authors:** Charlotte Bourdon, Jérôme Cachot, Patrice Gonzalez, Patrice Couture

## Abstract

Since the global ban on tributyltin in antifouling paints in 2008 by the International Maritime Organization, new products have been developed and brought to the market. Among them, copper pyrithione (CuPT) is used, but its mechanisms of toxicity remain little known. This project aimed to identify and measure the impacts of aqueous exposure to CuPT, an organic compound, and compare it to ionic Cu^2+^ added in the form of its inorganic salt CuSO_4_, in equivalent Cu^2+^ molar concentrations, on rainbow trout (*Oncorhynchus mykiss*) juveniles under controlled laboratory conditions. A 24-hour acute exposure was performed with nominal concentrations of 50 and 100 µg/L Cu from either CuSO_4_ or CuPT (labelled CuSO_4__50, CuSO_4__100, CuPT_50 and CuPT_100, respectively). The CuPT_100 condition induced 85 % mortality in 15 hours and the CuPT_50 condition induced 5 % mortality in the same period. A chronic exposure was then performed with nominal concentrations of 1 and 10 µg/L Cu from CuPT and 10 µg/L Cu^2+^ from CuSO_4_ (labelled CuSO_4__1, CuSO_4__10, CuPT_1 and CuPT_10, respectively). Measured aqueous concentrations of Cu^2+^ were slightly higher than nominal concentrations for the lower concentrations, but lower for the CuPT_10 condition. The 8- and 16-day toxicokinetics showed a greater accumulation of copper in the gills of fish exposed to CuPT compared to fish exposed to Cu^2+^ from CuSO_4_. The CuPT_10 condition induced 35 and 38 % mortality after 8 and 16 days of exposure, while no mortality was observed in the CuSO_4__10 condition. The growth of juveniles was not impacted during the 16 days of exposure for any condition. The activity of antioxidant enzymes (CAT, SOD, GPx) did not respond to exposure to either contaminant. The expression of genes involved in the antioxidant response (*sod1*, *sod2*, *gpx*), detoxification (*cyp1a*, *mt1x*, *mt2x*), Cu transport (*ctr1*, *ctr2*, *slc11a2*), energy metabolism (*AcoAc*, *cox*, 12S) and cell cycle regulation (*bax*) strongly decreased at Day 8 in the gills and at Day 16 in the liver of CuPT-exposed fish in comparison to controls at the same time point. This study clearly showed that the toxicity of Cu in the form of CuPT was much higher than that of ionic Cu from CuSO_4_ and provides new information on the compound that will be useful to develop regulations concerning its use and release in the aquatic environment.

**Credit author statement:** Charlotte Bourdon: Methodology, Validation, Investigation, Writing – Original Draft. Jérôme Cachot: Conceptualization, Methodology, Validation, Investigation, Writing – Original Draft, Supervision. Patrice Gonzalez: Validation, Investigation, Writing – Original Draft, Supervision, Funding acquisition. Patrice Couture: Conceptualization, Methodology, Validation, Investigation, Writing – Original Draft, Supervision, Project administration, Funding acquisition.

## 1. INTRODUCTION

Biofouling, *i.e.* the adhesion of organisms to any submerged surface, causes economic and environmental consequences. The increase in friction forces induces fuel overconsumption (up to 40 %) and increased maintenance costs (Champ, 2000). Vessels colonized by organisms can also be vectors for non-native species, and these can become invasive and destabilize an entire ecosystem. The application of antifouling paint helps to fight against this colonization. Since the banning of tributyltin (TBT), a widely used, highly effective but very toxic compound, at the end of the 20^th^ century, other copper (Cu)-based paints (Cu_2_O, CuCN) have been put on the market (Konstantinou and Albanis, 2004). One or more co-biocides are generally added to act on Cu-resistant organisms (Voulvoulis, 2006). Among them, copper pyrithione (CuPT), an organo-copper compound formed from two pyrithione ligands and a copper cation (Cu^2+^) in the centre, is commonly used for its antifungal and antimicrobial action (Okamura and Mieno, 2006), particularly in Japan (J-Check, 2021). Pyrithione acts as a copper ionophore, interacting non-specifically with the plasma membrane facilitating copper transport across the plasma membrane but also intracellular membranes (Reeder et al., 2011). This can result in increasing cellular levels of copper which could damage ion-sulfur clusters of proteins essential for cell metabolism and growth.

Since the first attempts to quantify metallic PTs (CuPT, ZnPT) in environmental matrices in 1999, there are few data on environmental concentrations because CuPT is photosensitive and has an estimated half-life of 7.1 ± 0.2 min in water (Maraldo and Dahllöf, 2004; Turley et al., 2000). The end-product of PT degradation is pyridine-2-sulfonic acid (PSA), which is far less toxic than the parent compounds (Turley, 2000). Nevertheless, Harino *et al*. (2007) reported a concentration of 2.2 µg·kg^−1^ dw in the sediment collected in a bay in Japan. In the bay of Toulon, France, Cu concentrations in the sediment have been reported between 5.8 and 864 mg·kg^−1^ dw (Tessier, 2011). The presence of CuPT in the environment could induce toxic effects on non-target species. Given the lack of information on its toxicokinetic and toxicodynamic properties, ecotoxicological studies are needed. Apart from toxicity studies on microalgae and crustaceans, very little research has focused on sublethal effects such as growth, reproduction, or the biochemical responses of organisms to CuPT exposure (Walker, 2006; Mochida et al., 2011; Mohamat-Yusuff et al., 2018). Few studies have reported on the toxicity of CuPT in fish embryos. In a recent study, Shin et al. (2022) compared the toxicity of CuPT and ZnPT on embryonic flounder and reported that the former had stronger effects than the latter on mortality, malformations and on the transcription levels of genes related to heart, nervous system and fin development. Almond and Trombetta (2016, 2017) also observed several developmental issues in zebrafish embryos exposed to CuPT.

The rainbow trout (*O. mykiss*) is a model species whose life cycle and physiology are well documented, is easy to rear in the laboratory, and is particularly sensitive to many contaminants (Le Bihanic et al., 2014; Weeks-Santos et al., 2019). A few studies on metal PT toxicity have been done on the rainbow trout (*O. mykiss*) (Okamura et al., 2002; Yamada, 2006). In our study, two experiments were performed on juvenile rainbow trout in the laboratory, during which the fish were exposed to either CuPT or CuSO_4_ in equivalent molar Cu concentrations. These exposures were aimed at characterizing the toxicity of an organo-Cu compound, CuPT, relative to ionic Cu^2+^ from CuSO_4_ in equivalent molar Cu concentrations, from phenotypic and molecular points of view. The first exposure lasted less than 24 h with high concentrations of 50 and 100 µg Cu^2+^·L^−1^ and induced rapid mortality. The second experiment exposed juveniles for 16 days to sublethal concentrations of Cu of 1 and 10 µg Cu^2+^·L^−1^. These concentrations were chosen based on the results of the first experiment. Metal contamination typically leads to an overproduction of reactive oxygen species (ROS) which must be regulated by the antioxidant defence system. Toxicokinetic and toxicodynamic properties of CuPT and CuSO_4_ were monitored by sampling fish at the beginning, halfway through the exposure, and at the end of the 16-day exposure in various tissues. Mortality and growth were the two phenotypic indicators, while from a molecular point of view, analyses of the activity of the antioxidant enzymes catalase (CAT), superoxide dismutase (SOD), and glutathione peroxidase (GPx) and analyses of gene expression levels were monitored. This study focused on the expression of 17 genes of interest. Genes selected included some involved in Cu transport (*ctr1* and *ctr2* (copper transporters 1 and 2) and *slc11a2* (solute carrier family 11 member 2 or divalent metal transporter)), antioxidant capacities (*gpx1* (glutathione peroxidase), *sod1* (cytoplasmic superoxide dismutase Cu/Zn), *sod2* (mitochondrial superoxide dismutase Mn) and *cat* (catalase)), detoxification (*cyp1a1* (cytochrome P450 family 1 subfamily A1), *gstA* (glutathione S-transferase A) and *mt1x* and *mt2x* (metallothionein isoforms 1X and 2X)), energy metabolism (*tgl* (triacylglycerol lipase-like), *cox1* (cytochrome c oxidase subunit 1) and *12s* (small mitochondrial ribosomal RNA)) and cell cycle regulation (*tp53* (cellular tumor protein tp53)). The objectives of this study were to (1) compare the toxicity threshold and the spectrum of sublethal effects after 8 and 16 days of exposure to CuPT or CuSO_4_ (2) compare the accumulation of Cu in tissues and (3) compare the mechanisms of toxicity of both compounds.

## 2. MATERIALS AND METHODS

### 2.1. Chemical preparation

Stock solutions of CuSO_4_ (39 mg/L) were prepared by dissolving CuSO_4_·5H_2_O (CAS# 7758-99-8; purchased from Sigma-Aldrich) in distilled water, and by serial dilutions of the stock solution. Stock solutions of CuPT (50 mg/L) were prepared in the dark by dissolving CuPT (CAS# 14915-37-8, purchased from Acros Organics, now Thermo Fisher Scientific) powder in the nontoxic organic solvent dimethyl sulfoxide (DMSO, CAS# 67-68-5) final concentration < 0.1 %) and distilled water, and by serial dilutions of the stock solution. Stock solutions were kept under dark conditions until use, to avoid photolysis.

### 2.2. Experimental conditions

Rainbow trout (*O. mykiss*) juveniles (9.9 ± 0.9 cm; 8.1 ± 1.6 g) were provided from pisciculture Saint-Alexis-des-Monts Inc. (Québec). Fish arrived at 7°C. A thermal acclimation was performed until the target temperature of 11°C was reached, at a rate of 1°C·day^−1^. Exposures were initiated after thermal and environmental acclimation, which lasted a total of six weeks. Acclimation and exposures were performed in reconstituted water ([Ca^2+^] 70 μM, [Cl^−^] 129 μM, [K^+^] 12 μM, [Mg^2+^] 13 μM, [Na^+^] 179 μM, [SO_4_^2-^] 63 μM). The first exposure included a control (0), and the molar equivalent concentration of ionic copper (Cu^2+^), at 50 and 100 µg Cu·L^−1^ from CuPT or CuSO_4_ to compare the range of lethal concentrations of both contaminants. There were 20 fish per tank, and one tank per condition. For the second exposure, 20 juveniles per tank were exposed during 16 days to the molar equivalent concentration of Cu^2+^, with 0 (control), 1 and 10 µg Cu·L^−1^ from CuPT (5 and 50 µg CuPT·L^−1^) or 10 µg Cu·L^−1^ from CuSO_4_ (40 µg CuSO_4_·L^−1^) to follow sublethal parameters. Every two days, three quarters of the exposure medium were renewed (static renewal experiments), and water quality parameters (temperature, nitrite, nitrate, ammonium, pH) and contamination were measured. Appropriate amounts of each stock solution were dispersed in each tank after water renewal to attain a designated nominal concentration for the exposure medium. Water samples were taken after each water renewal (beginning of the dark period to avoid photodegradation) and the morning after (during the light period). Both samples were analysed to monitor photodegradation. Water samples to quantify Cu were acidified with 0.2 (v/v) nitric acid 70 % Optima grade and kept in dark and cold at 4 °C until analysis. Water samples to quantify CuPT were kept in dark and cold at 4°C until analysis the same day. All conditions of the second experiment were in triplicate (12 tanks in total). Nominal concentrations of Cu were used to label the exposure conditions, and abbreviated as CuPT_1, CuPT_10, CuPT_50, CuPT_100, CuSO_4__10, CuSO_4__50 and CuSO_4__100. The experiments were carried out in oxygenated 40 L glass tanks in an environmentally controlled room (constant temperature at 11°C, light/dark cycle 14:10 h). Fish were fed daily *ad libitum* with pellets provided by the fish farmer (Nutra Fry®). Fish were checked daily for mortality and considered dead if they had no reactions after stimulation and if no movement of the mouth and the opercula could be detected. All procedures were approved by the INRS Animal care committee.

### 2.3. Fish sampling

The first experiment lasted less than 24 h. Dead fish were collected, and fish still alive were sacrificed. Tissues (liver, gills, and a sample of axial muscle collected above the lateral line and below the dorsal fin) of all fish were collected for Cu measurement. For the second experiment, fifteen fish were sampled at the start of the experiment (Day 0). Ten fish were also randomly sampled from all tanks (30 fish per condition) after 8 and 16 days. Fish sampled were sacrificed by a blow to the head and whole-body length and weight were recorded. Samples of gills, liver and muscle were collected to measure tissue Cu concentrations. These samples were frozen and stored at −20 °C until further analysis. Samples (20 mg) of liver and gills were kept in RNAlater® at −20 °C for genomic analyses. These two organs were chosen for transcriptomic analyses because gills is the main route of exposure and the liver is the central internal compartment for Cu accumulation and homeostasis (Grosell *et al*., 1998). Finally, liver samples (about 10 mg) were collected and immediately frozen in liquid nitrogen and stored at −80 °C for determination of antioxidant capacities. For the determination of Cu concentration, genomic and antioxidant capacity analyses, five fish per tank (15 per condition) were selected for analysis. The same individuals were used for all three analyses.

### 2.4. Cu and CuPT analysis

Fish tissue Cu analysis was performed by inductively coupled plasma-mass spectrometry (ICP-MS) (Model x-7, Thermo Elemental). Frozen tissues were lyophilized for 48 h (FTS Systems TMM, Kinetics Thermal Systems, Longueuil, QC, Canada), weighed, and then digested in 1 mL of nitric acid (70 %, v/v, Optima grade, Fisher Scientific) for 48 h. Then, 0.5 mL of hydrogen peroxide (30 %, v/v, Optima grade, Fisher Scientific) was added for an additional 48 h. Finally, ultrapure water was added to stop the digestion in a final digestion volume of 10 mL. Certified standards (DOLT-5 and TORT-3, n=5) were treated along with the fish tissue samples and allowed to estimate the efficiency of the digestion procedure and analytical accuracy. Recovery rates were 99% and 93%, respectively.

Copper in water samples was analysed with an ICP-AES (Varian Vista XP Axial CCD Simultaneous ICP-AES, Agilent Technologies) or ICP-MS (Model X-7, Thermo Elemental), depending on Cu concentration. CuPT in water samples was analysed by LC-MSMS (TSQ Quantum Access Thermo Scientific). The separation was carried out on an ACME-C18 100 mm x 2.1 mm x 3.0 µm column, with a column temperature of 40 °C, elution with 85 % methanol (0.1% formic acid) and 15% water (0.1% formic acid, 10 mM acetate) for 5 min, with a flow rate of 0.25 mL·min^−1^ and an injection volume of 10 µL. Samples and standards (Atrazine-D5) were diluted with the methanol/water solution using the ratio 85/15 (v/v).

### 2.5. Antioxidant capacities

Tissues samples were homogenized and crushed in a buffer solution prepared with 20 mM HEPES, 1 mM EDTA and 0.10 % Triton X-100. Aliquots were set aside for total protein determination by Lowry assay (Lowry *et al*., 1951). Enzyme activities included the quantification of catalase (CAT), superoxide dismutase (SOD) and glutathione peroxidase (GPx) activity. Analyses were performed using a UV/Vis spectrophotometer (Varian Cary 100, Varian Inc., Palo Alto, California, USA) on 96-well microplates at room temperature (20°C). Assay kits were purchased from Cayman Chemical Company Inc. (Ann Arbor, Michigan USA), and assays followed the manufacturer’s protocols. Catalase (kit No. 707002) activity was measured at 540 nm. Superoxide dismutase (kit No. 706002) activity was measured at 450 nm. Glutathione peroxidase (kit No. 703102) activity was measured at 340 nm. Enzyme activities are expressed as nmol/min/mg protein for both CAT and GPx and as U/mg protein for SOD (one unit of SOD is defined by the manufacturer as the amount of enzyme needed to exhibit 50 % dismutation of the superoxide radical). Protein concentrations were determined on liver homogenates before centrifugation using Coomassie (Bradford) Protein Assay Kit (No. 23200) at a wavelength of 595 nm and protein concentrations are expressed as mg protein per g of liver wet weight (Bradford, 1976).

### 2.6. RNA extraction and real-time qPCR

In this study, we measured the transcription level of 17 genes (*slc11a2*, *ctr1*, *ctr2*, *gpx1*, *sod1*, *sod2*, *cat*, *cyp1a*, *gstA*, *mt1x*, *mt2x*, *AcoAc*, *tgl*, *cox1*, 12S, t*p53* and *bax),* Accession number and specific primer pairs are shown in Table 1.

**Table 1:**
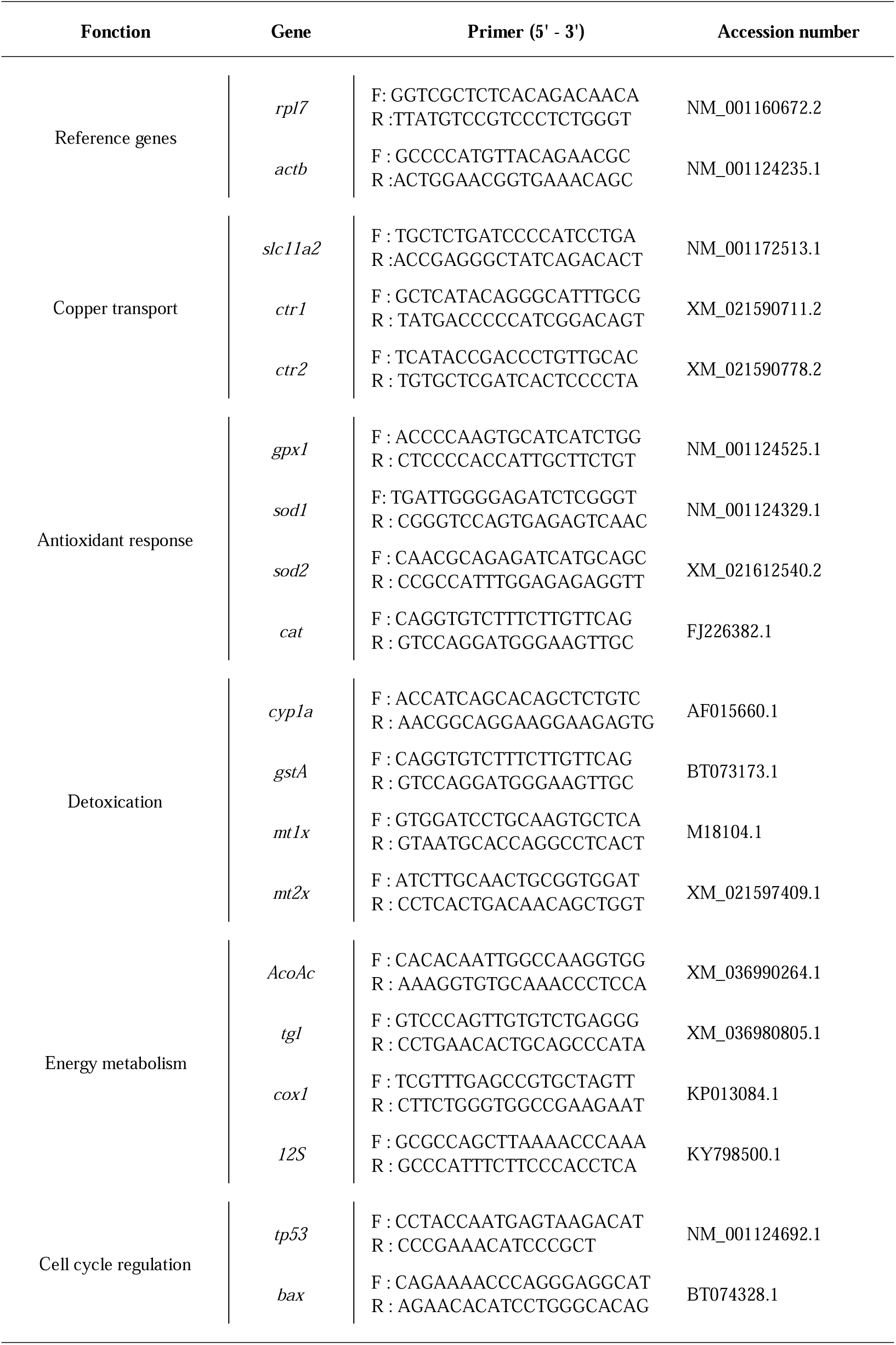
Accession number and specific primer pairs of 19 selected genes from the rainbow trout *O. mykiss*.

Gills and liver from 15 individuals per experimental condition were used for RNA extraction. Extraction was conducted with the kit Promega SV Total RNA Isolation System assay. Briefly, phenol-chloroform-isoamyl alcohol (25:24:1) was added to the homogenate, vigorously shaken and centrifuged (13 500 rpm for 5 min at room temperature). The upper aqueous layer (about 500 µL) was transferred to a new tube without disrupting the interface and one volume of ethanol 75% was added. DNA digestion was performed with the RNase-Free DNase I set for 15 min at 37°C. Reaction was stopped with the DNase Stop Solution and samples were washed with the RNA Wash Solution provided by the kit. RNase-free water was added to the spin column at the end of the process to eluate total RNA from the column. Total RNA quality and concentration for each extract were determined using absorbance measures at 260 and 280 nm with the spectrometer Epoch plate reader (Take3, Biotek) and analysed with the Gen5 software. The quality of RNA extracts with a 260/280 ratio upper than 1.8 was considered satisfactory.

Reverse transcription was conducted with the Promega GoScript Reverse Transcription System assay. One microliter of oligo dT at 1µM and hexa-primers at 1µM were added to 1 µg of total RNA with RNase-free water and incubated in thermocycler 5 min at 70°C then 5 min at 4°C. Then 8 µL of a mix (GoScript reaction buffer containing MgCl_2_, PCR Nucleotides, Recombinant RNasine Ribonuclease Inhibitor, GoScript Reverse Transcriptase) provided by the kit was added to start the reverse transcription reaction in the thermocycler (Eppendorf Flexide Mastercycler Nexus) for 5 min at 25°C, 1 h at 42°C and then kept at 4°C. Samples were kept at −20°C until real-time qPCR (rt qPCR). Rt qPCR was conducted with Promega GoTaqR qPCR Master Mix. Each well of the 384 wells plate has been filled with 3 µL of samples containing 30 ng of cDNA and 9 µL of master mix with SYBR green, reverse and forward primers pair at 2 µM each for one of the 17 selected genes and RNase-free water for a final volume of 12 µL. Real-time quantitative PCR (qPCR) analysis was performed using a LightCycler®480 (Roche), with a first cycle at 95°C for 2 min, followed by 45 cycles at 95°C for 15 sec and at 60°C for 1 min.. The reaction specificity was determined for each reaction from the dissociation curve of the PCR product. It was obtained by following the SyberGreen fluorescence level during a gradual heating of the PCR products from 60 to 95°C for 2 min. The relative quantification of each gene expression level was normalized to the arithmetic mean housekeeping genes *actb* and *rpl7*, and ΔCt values were recorded. From this comparison, fold-change factors were obtained for each gene by comparing each mean (n=15) value observed in the contaminated conditions with that of the corresponding control according to 2^−ΔΔCt^ method (Livak and Schmittgen, 2001).

### 2.7. Statistical analyses

Normality (Shapiro-Wilks) and variance homoscedasticity of residuals (Levene) were verified with a p-value set at 0.05. Since these assumptions of normality were not met, non-parametric tests were performed, as indicated in figure and table legends. Statistical analyses were performed using Excel’s statistical functions and Statistica. Differences between control and exposed conditions were considered significant if the p-value was below 0.05.

## 3. RESULTS

### 3.1. Cu concentration in the water

The Cu concentrations in water for the two compounds are shown in figure 1A. The concentration in the control aquaria was between 0.2 and 1.4 µg·L^−1^ (with an outlier of 3.3 µg·L^−1^). The measured concentrations in aquariums of the condition CuPT_1 were between 0.9 and 2.8 µg·L^−1^, with 50 % of the values between 1.3 and 2.2 µg·L^−1^. The measured concentrations in aquariums of the condition CuPT_10 were between 1.1 and 14.4 µg·L^−1^, with 50 % of the values between 2.4 and 4.4 µg·L^−1^. The measured concentrations in tanks of the condition CuSO_4__10 were between 1.3 and 12.2 µg·L^−1^, with 50 % of the values between 6.1 and 10.0 µg·L^−1^. Since there was a Cu background value in the controls, the exposure values of the CuPT_1 condition were higher than the nominal value. On the other hand, the exposure values of the CuPT_10 condition were strongly below the nominal value, with 50 % of the values between only 24 and 44 %, while the values of the CuSO_4__10 condition were quite close to the nominal value (50 % of the yield between 60 and 100 %).

**Figure 1:**
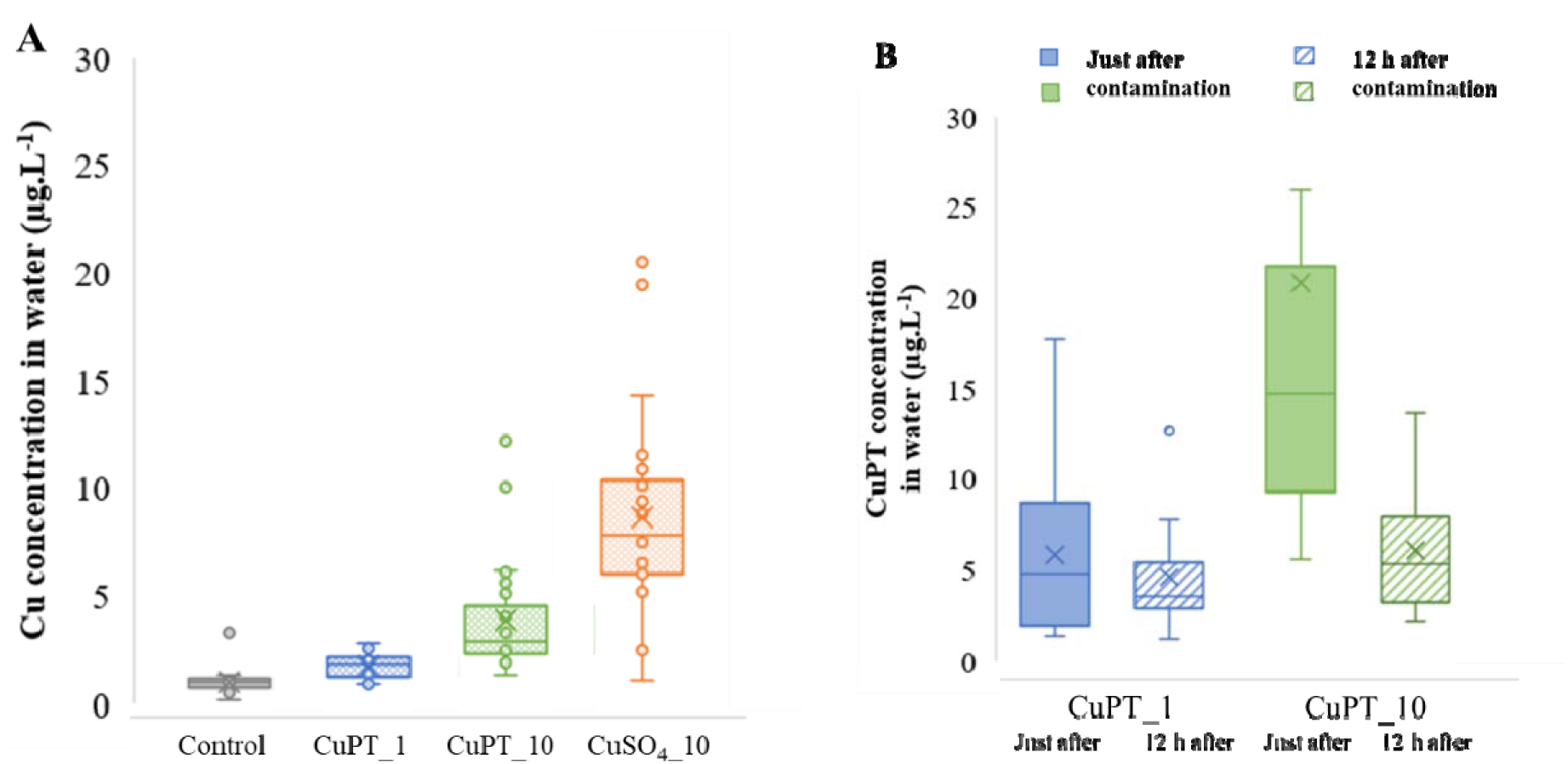
Boxplot of the gross values of Cu (A, n=15) and CuPT (B, n=10) concentration in water (µg·L^−1^) on the sampling days for the different conditions of exposure to CuPT and CuSO_4_. For Cu analyses: boxplot of all the values (just after contamination and 12 h after contamination, combined) for control (grey, n=24), CuPT_1 (blue, n=30), CuPT_10 (green, n=26) and CuSO_4_ (orange, n=26). For CuPT analyses: boxplot of the values just after contamination (n=13) and 12h after contamination (n=13), same colours for CuPT_1 (n=14) and CuPT_10 (n=14).

### 3.2. CuPT concentration in the water

The CuPT concentrations in water just after contamination, and 12 h after contamination for the two concentrations are shown in figure 1.B. The concentration in aquariums of condition CuPT_1 was between 2.7 and 12.0 µg·L^−1^ just after contamination with 50 % of the values between 3.3 and 8.0 µg· L^−1^, and between 2.5 and 14.0 µg· L^−1^ 12 h after contamination, with 50 % of the values between 4.2 and 5.8 µg· L^−1^. The concentration in aquariums of the condition CuPT_10 was between 6.9 and 27.2 µg· L^−1^ (outlier of 59.0 and 66.0 µg· L^−1^), just after contamination with 50 % of the values between 11.3 and 21.5 µg· L^−1^, and between 3.4 and 15.0 µg· L^−1^ 12 h after contamination, with 50 % of the values between 4.8 and 9.0 µg· L^−1^. The concentrations of CuPT obtained for the CuPT_1 condition coincide with the nominal concentration of CuPT (condition of 1 µg Cu· L^−1^ equals to 5.0 µg CuPT· L^−1^) while the values for the CuPT_10 condition are below the nominal concentration of CuPT (condition of 10 µg Cu· L^−1^ equal to 50.0 µg CuPT· L^−1^).

### 3.3. Cu accumulation in tissues

The mean Cu concentration in the liver of juveniles ranged from 165 ± 63 to 255 ± 58 µg·g^−1^ dw (figure 2A). No accumulation kinetics were visible between the beginning and the end of the exposure. Copper accumulation in liver was not compound-dependent. Only the fish in the CuPT_1 condition at Day 16 showed a significant increase of Cu accumulation in liver compared to the control. There was no significant difference in the liver of fish from the other conditions. In contrast, gill Cu accumulation varied among treatments (Figure 2B). Fish from the control and CuSO_4__10 conditions at the three days of sampling did not accumulate Cu in their gills (5 to 6 µg·g^−1^ dw). Conversely, fish from the CuPT_10 condition showed an accumulation of 141 ± 87 µg·g^−1^ dw on day 8 and 138 ± 49 µg· g^−1^ dw after 16 days. Gill Cu content in fish from the CuPT_1 condition did not differ from the controls. Remarkably, in fish exposed to the CuPT_50 and CuPT_100 conditions (first exposure), we observed a significantly greater accumulation of Cu in the gills than in the control after less than 24 h of exposure, with 72 ± 20 µg· g^−1^ dw for CuPT_50 and 68 ± 15 µg·g^−1^ dw for CuPT_100 (data not shown). Finally, the Cu contents in the muscle of fish exposed to CuPT_1, CuPT_10 or CuSO_4__10 for 8 or 16 days were all very low, between 1.8 and 2.7 µg·g^−1^ dw, and did not differ from their controls (figure 2.C).

**Figure 2:**
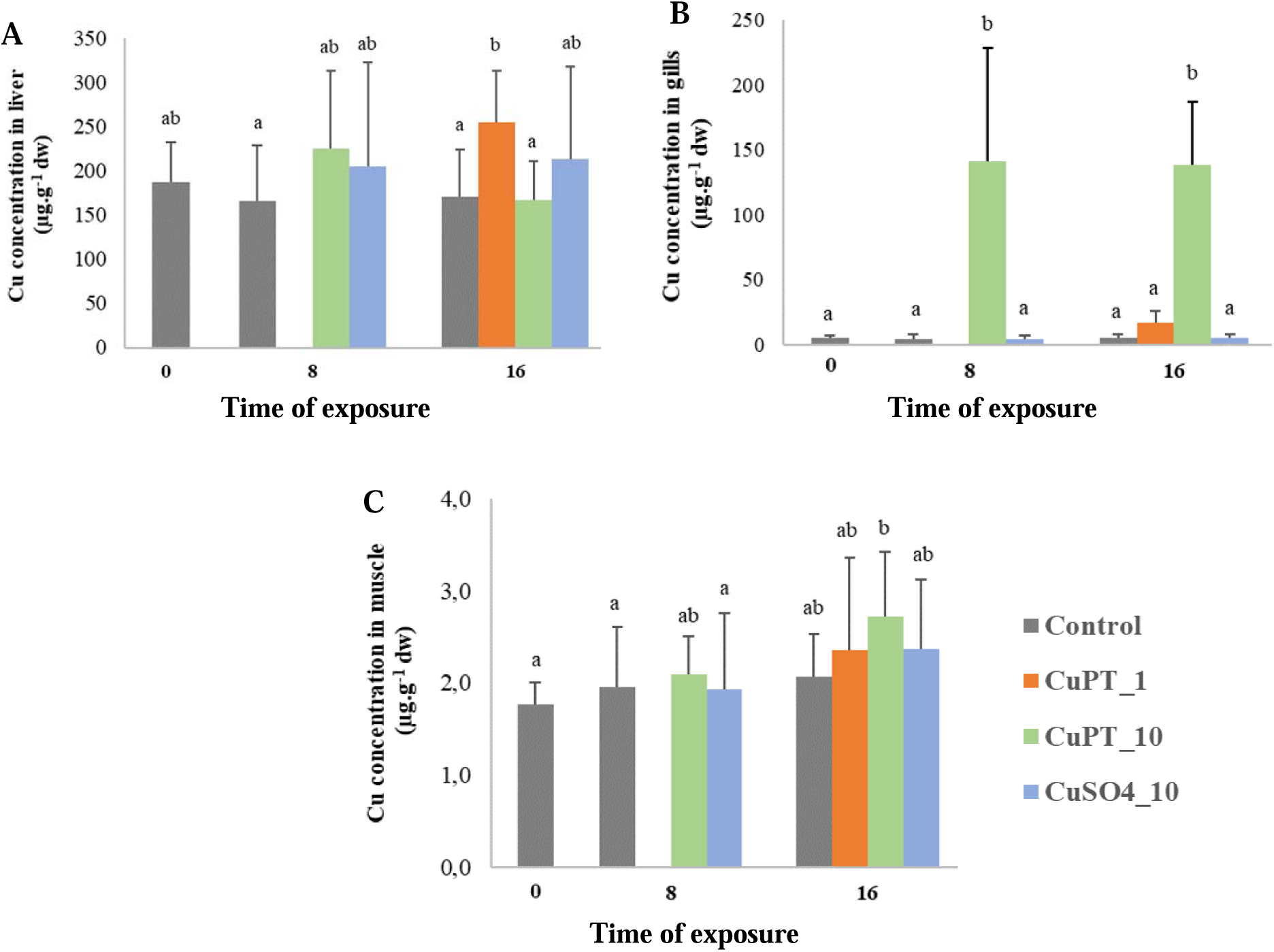
Cu concentrations in liver (A), gills (B) and muscle (C) according to the duration of the exposure to CuPT or CuSO_4_ (µg·g^−1^ dw) (n = 15, mean + SE). Different letters indicate a significant difference among conditions (Kruskal-Wallis test followed by Dunn’s test; p <0.05).

### 3.4. Mortality

During the daily water change, dead fish were counted and removed from the tanks. In the first experiment, there was 85 % mortality for the CuPT_100 and almost 5 % mortality for the CuPT_50 condition after less than 15 h of exposure (data not shown). No mortality was observed for the CuSO_4__50 or CuSO_4__100 condition during that period. The first experiment was stopped after these observations, which allowed to set the highest concentration for the second exposure (16-day chronic exposure) to 10 µg/L. For the second experiment, the percentage of cumulative mortality for CuPT_10 condition was significantly different from the control after 8 and 16 days of exposure with 38 % and 43 %. There was 20 % of mortality after 8 days of exposure to CuPT_1 and this value was maintained after 16 days. There was no mortality for control, and CuSO_4__10 conditions after 16 days of exposure (Figure 3).

**Figure 3:**
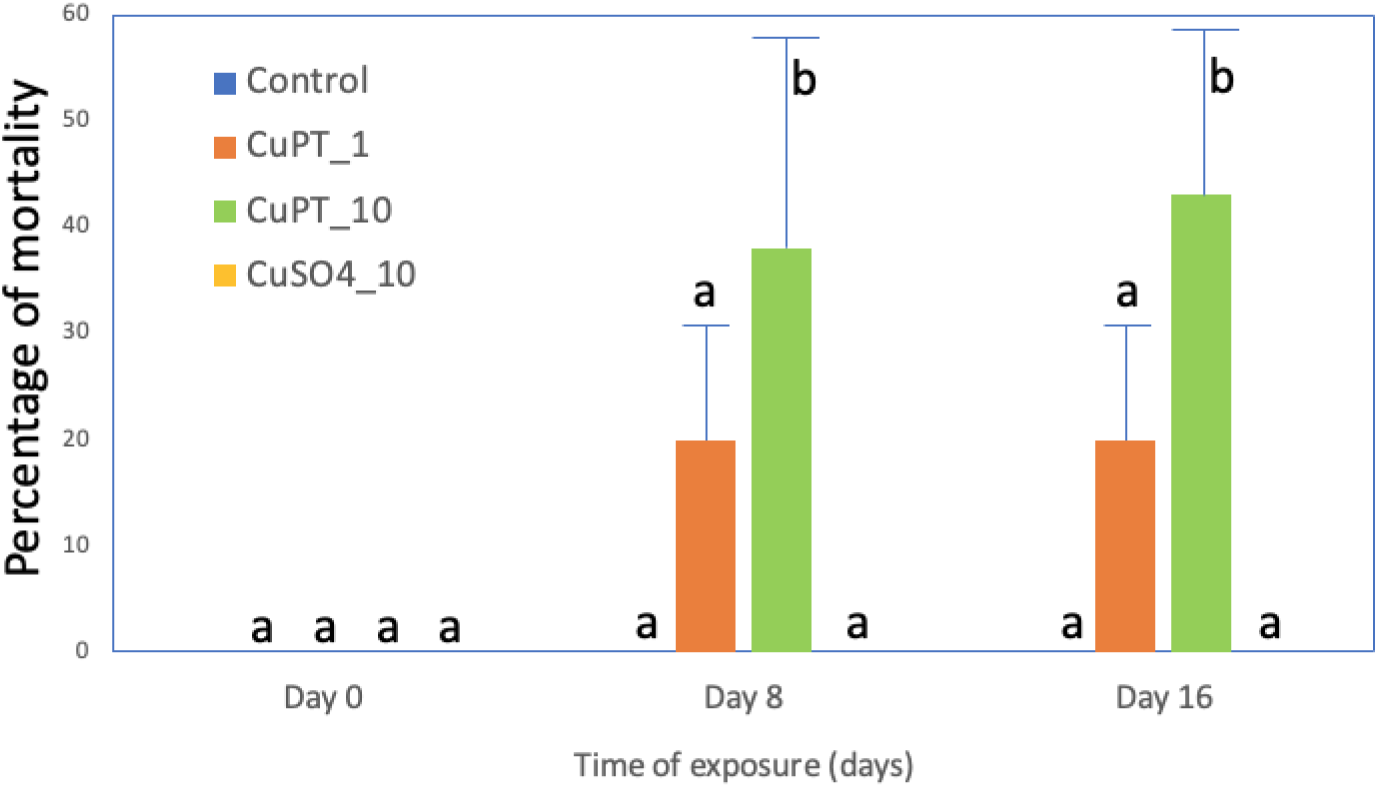
Percentage of mortality of juvenile depending on the time of exposure for all conditions (no mortality observed for control and CuSO_4__10; Dunnett’s test, p<0.05; mean +SE; n=3 aquariums per condition).

### 3.5. Biometric parameters

The fish sampled on Day 0 had an average length of 9.8 ± 0.9 cm and an average mass of 8.1 ± 1.6 g wet weight. After the 16 days of exposure, all fish had grown, reaching an average length of 10.2 ± 0.2 cm and an average mass of 10.4 ± 0.2 g wet weight. No significant difference in growth was observed among the conditions after the 16 days of exposure to either contaminant.

### 3.6. Antioxidant capacities

At the start of exposure (Day 0), mean CAT and SOD activities were 2562 ± 641 and 39 ± 11 nmol·min^−1^·mg^−1^ proteins, respectively. There was no significant difference in CAT and SOD activity among exposure days and exposure conditions (Figures 4A and B). The mean GPx activity on Day 0 was 37 ± 11 nmol·min^−1^·mg^−1^ proteins. The GPx activity was significantly reduced for all conditions on Day 8 and ranged from 18 ± 8 to 23 ± 10 nmol·min^−1^·mg^−1^ proteins. The mean GPx activity at Day 16 was not significantly different from the values at Day 0 and Day 8 (Figure 4C).

**Figure 4:**
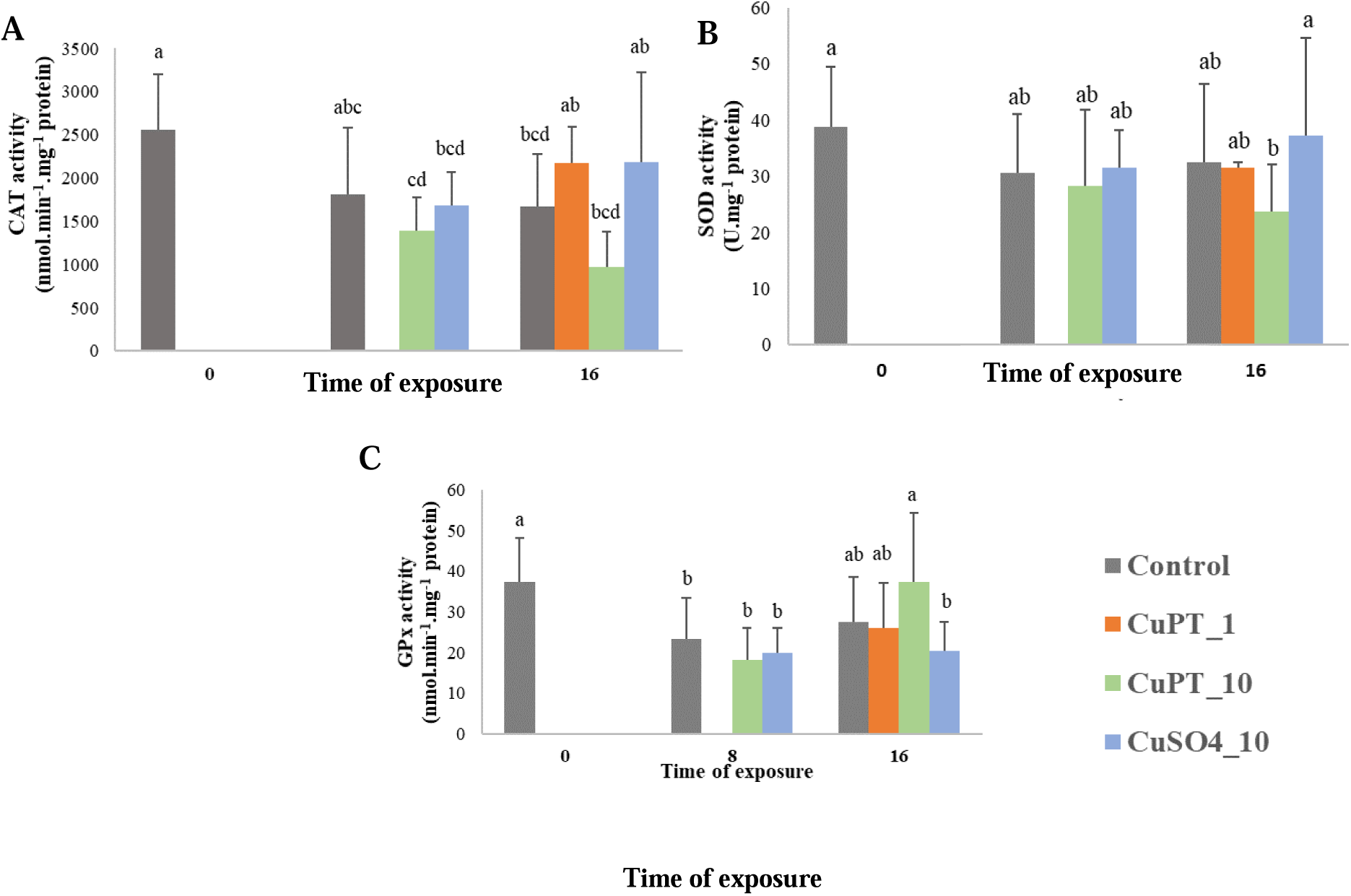
Enzyme activities in liver of rainbow trout in nmol·min^−1^·mg^−1^ proteins for CAT (A), SOD (B), and GPx (C) depending on the duration of exposure to CuPT or CuSO_4_ (n = 15, mean + SE). Different letters indicate a significant difference among conditions (Kruskal-Wallis followed by Mann-Whitney U test with Bonferroni correction, p <0.05), no data for CuPT_1 at Day 8.

### 3.7. Gene expression by real-time qPCR

After 8 and 16 days of exposure to CuPT and CuSO_4_, the expression of several genes varied significantly compared to the control (Table 2). There are no results for condition CuPT_1 at Day 8 because no fish were sampled. Genes *cat*, *gstA*, *tp53* and *tgl* showed no significant variation in their expression following exposure to the two contaminants, in either tissue (data not shown). CuPT_10 strongly repressed gene expression in gills at Day 8 for genes *sod1*, *sod2*, *gpx1*, *cyp1a*, *cox1*, 12S, *bax*, *ctr1*, *ctr2*. In contrast, several genes were overexpressed in the liver after the same exposure time in particular *gpx1*, *mt1x*, *mt2x*, *cox1*, *ctr1* and *AcoAc*. Also at Day 8, for CuSO_4__10 in liver, the gene expression profiles were different to those of fish exposed to CuPT_10. There was indeed overexpression of *mt1x* and *mt2x* (like CuPT_10), *ctr1*, but there was also a repression of *slc11a2* and *AcoAc*. In the gills of fish exposed to CuSO_4__10, there was overexpression of *mt1x* and *mt2x*, and repression of *gpx1*, *cox1*, *ctr1* and *ctr2*. After 16 days of exposure, the expression levels showed widely different trends than after 8 days. We observed a repression of all genes in the liver for all three exposure conditions CuPT_1, CuPT_10 and CuSO_4_ (except *ctr1* during the CuPT_10 exposure condition). In the gills, fewer genes responded compared to the liver. For CuSO_4__10, only one gene, *mt2x*, was repressed compared to the control. For CuPT_1, the *cyp1a* and *AcoAc* genes were repressed, and the *mt1x*, *mt2x* and *bax* genes were overexpressed. For CuPT_10, only the *cyp1a* gene was repressed, while the *mt1x*, *mt2x* and *ctr1* genes were overexpressed. Overall, our data show a rapid response in the gills following Cu contamination, with transcription levels of genes related to oxidative stress, detoxification and Cu transport functions altered, and these responses decreased at the end of the experiment. Conversely, the response was delayed in the liver, starting with overexpression of a few genes, then repression of most of the genes at the end of the exposure.

**Table 2:** Expression factor of genes of interest in gills and liver of juveniles of rainbow trout exposed to CuPT or CuSO_4_ (n=15). Only significantly different results from the control are shown (t test, p<0.05), overexpression and repression are indicated with + and – signs.

|  | Gills |  |  |  |  |  | Liver |  |  |  |  |
| --- | --- | --- | --- | --- | --- | --- | --- | --- | --- | --- | --- |
|  | Day 8 |  | Day 16 |  |  |  | Day 8 |  | Day 16 |  |  |
|  | CuPT_10 | CuSO<sub>4</sub>_10 | CuPT_1 | CuPT_10 | CuSO<sub>4</sub>_10 |  | CuPT_10 | CuSO<sub>4</sub>_10 | CuPT_1 | CuPT_10 | CuSO<sub>4</sub>_10 |
| *sod1* | -0.42 | / | / | / | / | *sod1* | / | / | -0.64 | -0.17 | / |
| *sod2* | -0.49 | / | / | / | / | *sod2* | / | / | -0.47 | -0.35 | -0.61 |
| *gpx* | -0.17 | -0.30 | / | / | / | *gpx* | +1.69 | / | -0.50 | -0.45 | -0.56 |
| *cyp1a* | -0.07 | / | -0.55 | -0.41 | / | *cyp1a* | / | / | -0.42 | -0.07 | / |
| *mt1x* | +3.09 | +1.38 | +2.32 | +13.42 | / | *mt1x* | +4.42 | +2.38 | -0.69 | / | -0.46 |
| *mt2x* | +8.06 | +1.77 | +2.16 | +16.64 | -0.73 | *mt2x* | +3.32 | +2.37 | -0.71 | / | -0.55 |
| *ctr1* | -0.60 | -0.68 | / | +1.45 | / | *ctr1* | +1.63 | +1.62 | -0.49 | +2.57 | / |
| *ctr2* | -0.33 | -0.77 | / | / | / | *ctr2* | / | / | -0.52 | -0.23 | / |
| *AcoAc* | +1.51 | / | -0.59 | / | / | *slc11a2* | -0.58 | -0.77 | -0.46 | / | / |
| *12s* | -0.18 | / | / | / | / | *AcoAc* | +2.49 | -0.43 | / | -0.32 | / |
| *cox* | -0.25 | -0.47 | / | / | / | *cox* | +1.51 | / | / | -0.31 | -0.47 |
| *bax* | -0.41 | / | +574 | / | / | *12s* | / | / | / | -0.21 | -0.42 |
|  |  |  |  |  |  | *bax* | / | / | -0.59 | -0.39 | -0.65 |

## 4. DISCUSSION

The objectives of this study were to (1) compare the toxicity and the spectrum of sublethal effects on rainbow trout juveniles after 8 and 16 days of exposure to CuPT or CuSO_4_; (2) compare the accumulation of Cu in tissues; and (3) compare the mechanisms of toxicity of both compounds by enzymatic biomarkers of antioxidant capacity and transcriptional response of selected genes.

### 4.1. Tissue accumulation of Cu and toxicity

The concentrations of Cu and CuPT in water were measured during the experiment and several phenomena were observable. Considering the process of photodegradation, the CuPT concentration measured just after tank contamination (in the dark) was at the desired targeted level following this contamination for the CuPT_1 condition (nominal CuPT concentration of 5.0 µg·L^−1^). The concentration remained close to target values 12 h later. The concentration of the CuPT_10 condition was lower than the nominal concentration (50.0 µg·L^−1^) just after the contamination and even lower 12 h later. The CuPT_1 condition met expectations of nominal Cu concentration. For the CuPT_10 condition, Cu concentrations did not reach 50 % of the nominal concentration. The CuSO_4__10 condition had Cu values very close to the nominal concentration. Therefore, juveniles exposed to CuPT_1 and CuSO_4__10 were exposed to the desired nominal concentrations, while those exposed to the CuPT_10 condition instead had exposure levels equivalent to 5 µg Cu^2+^·L^−1^. We hypothesize that CuPT had a strong tendency to adhere to inorganic and organic surfaces, such as aquarium and filter components and fish gills, a phenomenon that could be exacerbated at the higher concentration, leading to lower measured values than nominal aqueous concentrations.

Although the exposure to CuSO_4__10 corresponded to the targeted concentration, there was no Cu accumulation in the different tissues of the juveniles. Conversely, although fish from the CuPT_10 condition were exposed to lower than expected CuPT concentration, there was an accumulation of Cu in the gills, with significantly higher levels compared to the control and the CuSO_4__10 conditions. This difference in Cu accumulation between the two compounds imply that Cu is more bioavailable in the form of CuPT than in its ionic Cu^2+^ form from CuSO_4_. In their study, Borg and Trombetta (2010) showed that an exposure of juvenile brook trout (*Salvelinus fontinalis*) for 2 h at 16, 32 and 64 µg·L^−1^ of CuPT was sufficient to induce a significant accumulation of Cu in the gills. This very short exposure induced morphological modifications of the gills with the fusion of the secondary gill lamellae, induction of œdemas, loss of microridge structure and epithelial exfoliation. The authors observed the swelling of chloride cells and mitochondria and rupture of the lipid membranes. Another study reported severe damage to the gills of red sea bream (*Pagrus major*) after an exposure to CuPT (and ZnPT), including a dilation and fusion of secondary gill lamellae, necrosis and vacuolization as well as an expansion of the epithelial cells of the branchial cavity (Mochida et al., 2006).

In the liver of juveniles from our study, only the CuPT_1 condition induced a significant Cu accumulation after 16 days of exposure, while fish from the CuPT_10 and CuSO_4__10 conditions did not accumulate Cu in this organ. The expression of the genes involved in Cu transport (*ctr1*, *ctr2* and *slc11a2*) and detoxification (*mt1*, *mt2* and *cyp1a*) in the liver were all repressed for the CuPT_1 condition while the response was more moderate for the CuPT_10 and CuSO_4__10 conditions. These differential gene expressions, if they are accompanied by a decrease in the quantity of the corresponding proteins and of their activities, could explain the difference in Cu accumulation in the liver. Following waterborne exposure, teleost fish normally accumulate Cu in the gills (the main route of exposure) and the liver is the central internal compartment for Cu accumulation and homeostasis (Grosell *et al*., 1998). In juvenile rainbow trout, an exposure of 10 days to 20 and 100 µg·L^−1^ of CuSO_4_ induced accumulation of Cu in the gills but not in the liver, which does not coincide with the observations of our study (Shaw *et al*., 2012). Muscle is not a target organ for Cu storage, so it is not surprising that none of the exposure conditions studied led to an accumulation. Due to the low solubility of CuPT in water, it is expected to adsorb to suspended matter and food pellets. It would have been interesting to measure the Cu concentration in the digestive tract of our juveniles, to examine a potential accumulation of Cu by gut. Indeed, the relative efficiency of Cu uptake from food appears to be similar to the efficiency of Cu uptake from water filtered by the gills (Clearwater et al., 2002). In addition, it might have been interesting to follow the Cu levels in the tissues during a depuration period. These observations would have made it possible to know whether CuPT is rapidly eliminated from the gills, or whether it is instead transferred to the liver or metabolized and cleared from the body.

Only the CuPT_10 condition induced mortality after 8 and 16 days of exposure, while for CuSO_4__10 no mortality was observed during the 16 days of the exposure. Cu toxicity has been studied in several fish species. It is acutely toxic to rainbow trout with a 96-h LC_50_ of 210 µg·L^−1^ (De Boeck *et al*., 2004). This result could explain the absence of mortality in the second experiment with CuSO_4_ since the exposure concentration was 10 µg·L^−1^. The difference of mortality between CuPT and CuSO_4_ can be related to differences in the levels of Cu accumulation in the tissues. The bioavailability of Cu for the fish exposed to CuPT directly impacted their survival. This observation is strongly supported by the results of the first experiment, where in less than 24 h, the gills of the juveniles of the CuPT_100 condition had already accumulated 68 ± 15 µg·g^−1^ dw of Cu (compared to 5.53 ± 1.9 µg·g^−1^ dw for the controls) which had induced 85 % of mortality. In contrast to mortality, growth was not affected in our study under any experimental condition over the 16 days of exposure. We can assume that the growth of the juveniles did not have time to be impacted by the contaminants in 16 days compared to the control conditions. Our experimental design does not allow to calculate the lethal concentration 50 % of CuPT (LC50) after 8 or 16 days. In the literature, several studies have focused on the acute toxicity of CuPT, mainly on microalgae and crustaceans. Among these studies, growth inhibition after 72-h (72-h EC_50_) for the microalgae *Dunaliella tertiolecta* was 7.3 µg·L^−1^, while the values for the microalgae *Tetraselmis tetrathele* was 12 µg·L^−1^, for the microalgae *Chaetoceros calcitrans* was 3.2 µg·L^−1^ and 1.5 µg·L^−1^ for the diatom *Skeletonema costatum* (Onduka et al., 2010). Crustacean mortality after 24 to 96 h of exposure to CuPT has been studied on several species. Values reported were 830 µg·L^−1^ (24-h LC_50_) for the artemia *Artemia salina* (Koutsaftis and Aoyama, 2007) and 250 µg·L^−1^ (48-h LC_50_) for the same species (Lavtizar et al., 2018). Data for the *Tigriopus japonicus* copepod give 24-h LC_50_ = 41 µg·L^−1^ (after Yamada 2006) and 96-h LC_50_ = 30 and 32.7 µg·L^−1^ (Bao et al., 2014, 2011). Studies on the acute toxicity of CuPT on fish give 96-h LC_50_ = 7.67 and 9.3 µg·L^−1^ on *Pagrus major* (from Yamada, 2006; Mochida *et al*., 2006) and 96-h LC_50_ = 4.3 and 2.6 µg·L^−1^ on the fathead minnow *Pimephales promelas* (from Yamada 2006; Regulation (EU) No. 528/2012, 2014). Finally, Okamura *et al*. (2002) carried out a toxicity test on rainbow trout larvae (24 h post-hatch) with CuPT and other biocides (ZnPT, Irgarol 1051, diuron, Sea-Nine 211) for 28 days, at concentrations of 0, 1.0, 2.0, 4.0, 8.0, 16 µg·L^−1^. The order of toxicity of the compounds (based on nominal concentrations) on the trout larvae after 28 days of exposure was CuPT> ZnPT> Sea-Nine 211> KH101> diuron> Irgarol 1051. The authors evaluated 4 values of LC_50_, at 7, 14, 21 and 28 days, which gives respectively for CuPT exposure 7.6; 3.0; 1.7 and 1.3 µg·L^−1^, *i.e.,* 1000 times more toxic than for Irgarol 1051 and diuron. On marine medaka *Oryzias melastigma* larvae, the 96-h LC_50_ was reported to be 8.2 µg·L^−1^ (Bao et al., 2011). All these studies show to what extent aquatic species at all stages of life are sensitive to CuPT. In our study, juvenile rainbow trout were more tolerant than the aquatic species cited above. It could be interesting to carry out the same experiment on the larval stage of rainbow trout to have a comparison of these two life stages.

### 4.2. Oxidative stress and molecular responses

Free Cu generates hydroxyl radicals which are the source of ROS. To fight against ROS, antioxidant molecules (ascorbic acid, glutathione) and enzymes (SOD, GPx, CAT, etc.) can eliminate them. When ROS levels increase beyond the antioxidant capacities of the cells, this leads to oxidative stress. The 16-days exposure to CuPT and CuSO_4_ did not induce an increase in the activity of antioxidant enzymes, suggesting an absence of oxidative stress in the liver of these juvenile fish. Sanchez *et al*. (2005) showed a significant increase in SOD and CAT activities after 4 days of exposure to CuSO_4_ on the three-spine stickleback, at a concentration of 25 µg·L^−1^, then it returned to baseline after 8 days. Regarding GPx activity, only the 200 µg·L^−1^ condition induced an increase after 12 days of exposure in the same study. If, in our study, juveniles were sampled after 4 days of exposure, we may have observed a response of these antioxidant enzymes like that of Sanchez *et al*. (2005). Borg and Trombetta (2010) have shown that TBARS levels in the brook trout *Salvelinus fontinalis* gills were significantly increased following CuPT exposure conditions at 16, 32 and 64 µg·L^−1^. In parallel, there was a significant decrease of 10 and 25 % respectively in total antioxidant capacity (TAC) for conditions 32 and 64 µg·L^−1^ (Borg and Trombetta, 2010).

Given the absence of response at the level of antioxidant enzyme activity, the levels of gene expression observed in the liver provide interesting insights. Like for the CAT enzyme activity, there was no response from the *cat* gene transcription over the entire exposure period. The *sod1* gene did not show differential expression on Day 8, whereas on Day 16 it was significantly less expressed in the conditions CuPT_1 and CuPT_10 but was not deregulated in the condition CuSO_4__10. The *sod2* gene was repressed for all exposure conditions on the 16^th^ day of exposure. The *gpx1* gene was overexpressed only in the CuPT_10 condition, in contrast to the GPx enzyme which did not vary in comparison to the control and the other conditions. On day 16, the three exposure conditions induced a decrease in the expression of the *gpx 1* gene in contrast to the activity of the corresponding enzyme which did not vary. The lack of induction of enzymes involved in the response to oxidative stress could have been counterbalanced by the expression of metallothioneins. Indeed, these proteins have been described as being involved in the response to oxidative stress in a number of organisms (Ruttkay-Nedecky et al., 2013). Thus, the overexpression of *mt* genes probably contributed to the control of oxidative stress and limited the changes in antioxidant enzyme levels. The *nfe2.1* gene (nuclear factor erythroid 2 related factor 1) is a precursor of oxidative stress response and would have been interesting to study in addition to the *sod1*, *sod2*, *cat*, *gst* and *gpx1* genes to better understand the extent of oxidative stress that our exposure conditions induced.

The levels of gene expression show differential responses among tissues and conditions, and over time. In the gills, gene expression was predominantly repressed at Day 8 for the CuPT_10 condition, while in fish exposed to CuSO_4__10 fewer gene expression levels were altered. For both conditions, the antioxidant response genes and Cu transport genes were repressed, while the *mt1x* and *mt2x* genes were overexpressed. These early responses for CuPT_10 coincided with Cu accumulation in the gills, while molecular responses still occurred despite the absence of accumulation of Cu in the gills for the CuSO_4_ condition. Still in the gills, on Day 16, the levels of gene expression returned to the baseline for the three exposure conditions, except for the detoxification genes (*mt1x*, *mt2x*, and *cyp1a*) and pro-apoptotic gene (*bax*).

In juvenile trout, differential mechanisms between the absorption and metabolism of Cu are induced between the two Cu species. Indeed, there was an increase in the levels of transcription of the *mt1x* and *mt2x* genes induced by the two compounds, yet the accumulation of Cu in the gills was observed only for the conditions exposed to CuPT. Metallothioneins (MTs) are cysteine-rich proteins involved in maintaining sufficient intracellular supplies of certain essential metals such as Cu and Zn and detoxifying excess intracellular metals. The overexpression of these genes clearly shows that Cu has induced a molecular response (Amiard *et al*., 2006). Exposure to CuPT, but not to CuSO_4_, induced a repression of the *cyp1a* gene in the gills and the liver. Cytochromes P450 are a multigene family of heme-containing proteins that oxidize, hydrolyse, or reduce hydrophobic chemicals by inserting an oxygen atom to the substrates during the reaction cycle to increase their water solubility. They are present mainly in the liver of fish, but also in their gills and digestive tract (Varanasi 1989). In our study, detoxification seems to have been managed mainly by MTs rather than by cytochromes P450.

Regarding the regulation of the cell cycle, *p53* gene expression was not modified. On the other hand, the *bax* gene for CuPT_1 at day 16 was particularly overexpressed (+574), suggesting a peak of gill cell apoptosis at the end of the experiment. We hypothesise that the gills were initially the target organ, but that they were able to adapt by setting up molecular defence mechanisms against this contamination at the start of exposure. However, the significant increase in the *bax* gene after 16 days seems to indicate cellular damage in this organ at the end of the experiment. In the liver, the molecular response was quite different. There were fewer genes with an altered level of expression following exposure to our experimental conditions. The CuPT_10 condition predominantly caused overexpression of genes for detoxification, oxidative stress, and energy metabolism. At the same time, the CuSO_4__10 condition instead induced the repression of the Cu transport genes and an overexpression of the detoxification genes. Despite a lack of quantifiable accumulation of Cu in the liver, it still had measurable responses for both contaminants at the molecular level. Finally, on Day 16, almost all genes were differentially expressed under all exposure conditions and were mostly repressed. Our data indicate that in the liver, there was probably a massive cytotoxicity after 16 days of exposure, and that juveniles were no longer able to defend themselves against contamination, for both contaminants studied.

## 5. CONCLUSIONS

This study supports the greater toxicity of CuPT compared to CuSO_4_, for equivalent Cu concentrations, on juvenile rainbow trout. The major phenotypic response that we observed was the mortality of juveniles exposed to CuPT_10 (35 and 38 % on Days 8 and 16), CuPT_50 (5% in 15 h) and CuPT_100 (85 % in 15 h), while no mortality was observed for CuSO_4_ exposures up to 10 µg·L^−1^. The 16-days exposure to CuPT and CuSO_4_ did not affect the growth of the juveniles. Our study indicates that a concentration range of 1 to 100 µg Cu^2+^·L^−1^ of CuPT is toxic to juvenile rainbow trout while this is not the case for CuSO_4_. The higher toxicity of CuPT could be explained by the higher bioavailability of Cu in CuPT compared to CuSO_4_, as supported by the strong and rapid accumulation of Cu in the gills of fish exposed to CuPT. The activities of antioxidant enzymes (CAT, SOD, GPx) were not significantly altered, making it difficult to conclude on the oxidative stress generated by our exposures. Nevertheless, gene expression analyses showed the adaptive responses of juveniles to CuPT and CuSO_4_ in the gills, while the liver experiences cytotoxic effects at the end of the exposure. The mechanisms of action of CuPT have yet to be investigated through additional studies. Our study confirms the toxicity of CuPT in antifouling paints for juveniles of rainbow trout, a non-target species. This is particularly worrying since CuPT will naturally adsorb to suspended particles and settle to the sediment. Accumulation of CuPT in the sediment can impact species with at least one benthic life stage, which is the case for rainbow trout embryos and larvae. Further studies are clearly needed to evaluate the toxicity and the risk of CuPT on early life stages of fish.

## 6. AKNOWLEDGEMENTS

The authors express their gratitude to J. Perreault and J.F. Dutil for performing the quantification of Cu and in waters and in juvenile tissue, and S. Prémont and S. Moïse for the development of an analytical method to quantify CuPT in water. Also, we thank the instrumental platform of Molecular Biology for the formation to extraction and RT-qPCR (EPOC Laboratory, University of Bordeaux-Arcachon). This study was funded by a Discovery grant from the Natural Sciences and Engineering Research Council of Canada to P. Couture and by funds from the EPOC laboratory. The authors also wish to thank an anonymous reviewer who provided a critical review of an earlier version of this paper.

